# Crystalline guanine packed within vacuoles serves as nitrogen store in *Chromera velia*

**DOI:** 10.64898/2026.01.31.703024

**Authors:** Vijaya Geetha Gonepogu, Jana Pilátová, Dorsaf Ennaceur, Aleš Tomčala, Marie Vancová, Jitka Richtová, Robyn Roth, Ursula Goodenough, Miroslav Oborník, Peter Mojzeš, Ansgar Gruber

## Abstract

Nitrogen is an important element for all living organisms. Photoautotrophic organisms need to assimilate nitrogen from the environment, therefore changes in nitrogen availability have a strong influence on their growth and metabolism. Many microalgae have been known to contain crystalline inclusions, and recently, it has been shown that many of these consist of purines like guanine and thus must be linked to the cellular nitrogen metabolisms. The alveolate alga *Chromera velia* contains such guanine crystals, and during its lifecycle, the alga is thought to be subjected to strong changes in external nitrogen availability. Here, we investigated the formation or decline of crystalline guanine in dependence of the availability of inorganic nitrogen in the growth medium. Cells were examined using polarised light microscopy, Raman micro-spectroscopy, chromatography (HPLC), transmission and scanning electron microscopy. The cellular guanine crystal content decreased during nitrogen starvation and increased upon transfer of the cells back to standard growth medium containing nitrate. Raman micro-spectroscopy showed that the crystals were composed of anhydrous guanine in beta-polytype. They appear in unspecific positions throughout the cell, and staining with the green dye Lysotracker DND-26 suggests that they are within vacuoles. Stacks of crystals could be observed in cells via freeze fracture and freeze etching electron microscopy, which unambiguously showed a membrane around the crystal aggregates, in a similar arrangement as has been shown for guanine storage vacuoles (GSV) in *Chlamydomonas reinhardtii*. We developed a method to isolate the guanine crystals from whole cells, and were able to obtain crystals which retained their flat, plate-like structure, matching the electron microscopic observations from whole cells. The isolated crystals were shown to consist of nitrogen rich compounds via energy-dispersive X-ray (EDX) analysis, and Raman micro-spectroscopy confirmed that they consist of guanine.

## Introduction

Photosynthetic organisms, including algae and cyanobacteria, can fix carbon through photosynthesis. However, in addition to this ability, phototrophs rely on external nutrients, especially nitrogen and phosphorus, which are essential for cellular functions. Nitrogen, as a component of nucleic acids and proteins, is a limiting element for biological productivity and plays an important role in marine biogeochemistry (Gruber, 2008). In particular, the ratio between carbon, nitrogen and phosphorus (C:N:P), also known as the Redfield ration, is empirically found to be close to 106:16:1 (Redfield 1934). The Redfield ratio serves as a standard for understanding nutrient dynamics in marine ecosystems (Singh et al., 2015) or to predict the growth of phytoplankton (Arteaga et al., 2014; Takahashi et al., 1985). Changes in nutrient availability, such as nitrogen and phosphorus, however, can cause shifts in the Redfield ratio, pointing to flexibility in the storage compounds. While carbon storage for example in the form of starch or lipids is rather well investigated, there is increasing evidence that also nitrogen and phosphate can be accumulated and stored in the cell beyond immediate demand.

Most of the inorganic nitrogen in the ocean is present in the form of nitrate. The cells absorb nitrate, which is then converted into ammonia via nitrite and assimilated for the production of amino acids and other compounds (e.g. nucleic acids). Many photosynthetic eukaryotes can also utilize organic nitrogen sources such as urea, amino acids or purine bases such as guanine or uric acid. Nitrogen deficiency leads to numerous changes in the metabolism and cell biology of algae. These include an interruption in cell division, a slowdown in the photosynthesis and an accumulation of carbon (Sasaki et al., 2023). In recent years, the multifunctionality of crystalline guanine as a biological compound has been increasingly recognised (Mojzeš et al., 2020). Guanine is a purine, one of the four fundamental bases in nucleotides, that serve as integral components of nucleic acids (Gur et al., 2016). Guanine crystals are biogenic crystalline inclusions that have been identified in various eukaryotes (Pilátová et al., 2022; Sasaki et al., 2023). They are found in three structural forms: guanine monohydrate (Thewalt et al., 1971) and alpha- and beta polytypes of anhydrous guanine (Guille & Clegg, 2006; Hirsch et al., 2015). Anhydrous guanine crystals serve as a reservoir to store nitrogen that is used under nitrogen-deficient conditions (Moudříková et al., 2016) and are mainly found in marine and freshwater algae (Jantschke et al., 2019; Mojzeš et al., 2020; Goodenough et al. 2025). Monohydrate guanine crystals have also been identified in the bacterium *Aeromonas salmonicida* subsp. *Pectinolytica* and in marine diplonemids, heterotrophic protists related to euglenids, symbiontids and kinetoplastids (Pilátová et al., 2022; Sasaki et al., 2023; Tashyreva et al., 2022). Animals such as fish, reptiles and spiders possess skin and visual organs that exhibit a reflectance effect due to the presence of a structural color caused by their remarkably high refractive index (Jantschke et al., 2019), a property that is exploited in bio-optical systems (Aizen et al., 2018).

*Chromera velia,* an apicomplexan alga (Apicomplexa, Apicomonada; after Cavalier-Smith 2018) associated with corals, was first isolated from the scleractinian coral *Plesiastrea versipora* in Sydney Harbour (Moore et al., 2008). *Chromera velia* is phylogenetically closely related to apicomplexan parasites (Apicomplexa, Sporozoa) such as *Plasmodium* and *Toxoplasma,* which are known to cause serious diseases in humans and other animals (Oborník et al., 2011). *Chromera velia* exhibits a life cycle consisting of immotile, coccoid cells that subsequently transform into zoosporangia with motile flagellated zoospores (Oborník et al., 2011; Weatherby et al., 2011). *Chromera velia* is thought to infect coral larvae as an intracellular symbiont (photoparasite), which is a mixotroph that combines phototrophy and parasitism (Oborník 2020). As such, it is intended to alter a nitrogen-rich and nitrogen-pure environment and thus counteract the need to store biologically accessible nitrogen in the cell.

In this study, we demonstrated the presence of anhydrous beta-polytype guanine crystals in *C. velia* within cytosolic vacuoles, that are formed or consumed in response to inorganic nitrogen availability and therefore likely serve as nitrogen storage compartments in *C. velia.* The morphology of and cellular arrangement of the crystals was investigated in vivo and in vitro using polarised light microscopy and advanced microscopic techniques, such as SEM, TEM, and cryo-SEM in combination with Raman micro-spectroscopy.

## Material and Methods

### Culture conditions

All *Chromera velia* cultures were kept under artificial light, with a photo period of 12 h light/12 h dark, in 150 ml culture flasks. The light intensity was maintained at 30–50 μmol m^−2^ s^−1^ and a temperature of 26 °C. For the nitrogen availability experiment shown in Figure 1, 10 ml of a 7 day old stationary culture with an optical density of 0.1210 (OD_600nm_, measured with a TECAN Infinite 200 PRO spectrophotometer) was added to each flask as inoculum, triplicates were prepared for nitrogen-repleted (F/2 media) and nitrogen-depleted conditions (F2 media without nitrate). The average number of crystals per cell was calculated from day 0 to day 5. After day 5, the cultures grown under nitrogen-depleted conditions were centrifuged, and the pellet was washed twice with nitrogen-replete media and then transferred to nitrogen-replete conditions (Figure 1). Growth of the cultures was assessed via optical density (OD_600nm_, TECAN Infinite 200 PRO spectrophotometer).

**Figure 1:**
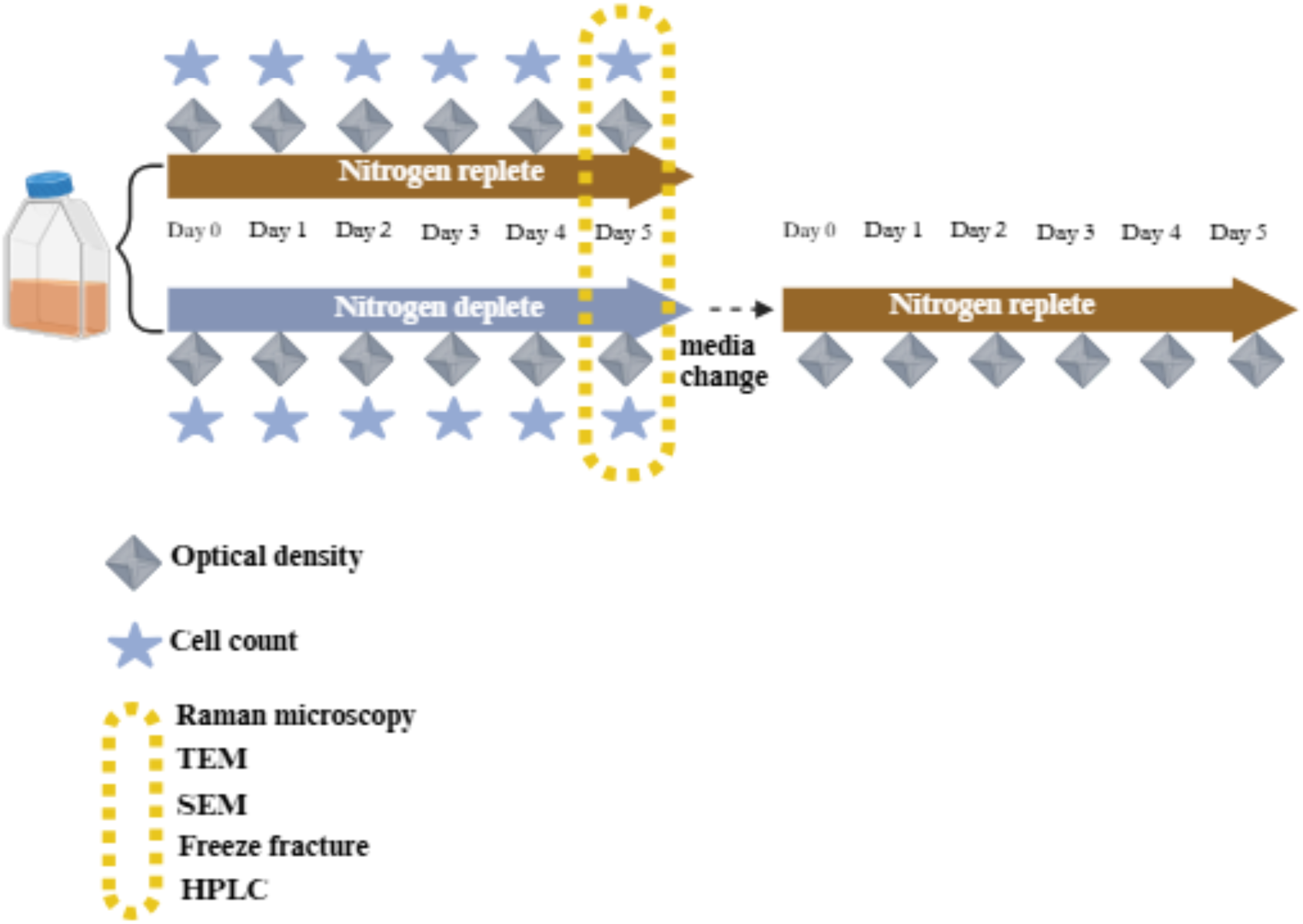
Experimental design for testing guanine crystal formation in response to nitrogen availability in *Chromera velia.* Starter cultures were inoculated in F/2 medium for 7 days (not shown). The cultures were then split and transferred to two conditions; nitrogen repleted (i.e. standard F/2 containing nitrate as an inorganic nitrogen source plus phosphates, trace metals and vitamins), and nitrogen depleted. Except for nitrate, the nitrogen-depleted media had the same composition as the standard F/2 medium. After day 7, *C. velia* cells cultivated in nitrogen depleted cultures were centrifuged and resuspended into nitrogen replete media, to observe the recovery from the nitrogen starvation.

### Light microscopy observations and vacuole staining

The cultures of *Chromera velia* under nitrogen-replete and depleted conditions were subjected to polarized light microscopy (Olympus DP20 Model BX53, Shinjuku, Tokyo, Japan) for a period of 5 days to observe the formation of crystals. Crystals in the zoospores of *C. velia* were identified according to the method described in Richtová et al (2023) and fixed in 4% paraformaldehyde under nitrogen-enriched conditions. For staining the vacuoles, labelling was performed on live *C. velia* cells in nitrogen-repleted conditions (5 days old), which were pre-treated with 0.1% Triton X-100 for 30 minutes prior to staining, for permeabilization. For optimal results, cells were incubated at a concentration of 10 µM in PBS buffer for 2 hours at 37°C. As negative controls, cells were treated only with either 0.1% Triton-X100 or 1% Dimethyl sulfoxide (DMSO) for the same incubation time and temperature. A fluorescence microscope the GFP filter set was used to image the cells, allowing the excitation of 51% of the probe (480 nm) with emission from 495 to 540 nm. The YFP filter set, which allows excitation wavelengths between 540 and 585 nm, also detects autofluorescence at 640 to 720 nm range.

### Raman analyses of Chromera velia cells under nitrogen-depleted and nitrogen-replete conditions

For *in-situ* determination of the chemical composition of intracellular structures, a confocal Raman microscope (alpha300 RSA; Oxford Instruments - WITec, Germany) was used as previously described (Barcytė et al., 2020; Pilátová et al., 2022). To immobilise the *C. velia* cultures on the quartz slide, 5 mL of the cell pellet of the nitrogen-repleted sample was mixed with 5 mL of 1% (wt/vol) solution of low-temperature-melting agarose (Carl Roth, Cat. No. 6351.5; Germany) in artificial sea water, immediately spread as a single-cell layer between a quartz slide and a coverslip, and sealed with CoverGrip sealant (Biotium, USA). A similar procedure was carried out for the nitrogen-depleted samples. Two-dimensional Raman maps were created using laser excitation at 532 nm (20 mW power in the focal plane) and the UPlanSApo 60×, NA 1.20 (Olympus, Japan) water-immersion objective. A scanning step size of 200 nm in both directions and an integration time of 100 ms per voxel were used. At least 30 cells were measured for both nitrogen-depleted and nitrogen-repleted cultures. The Raman spectra of the crystalline particles isolated from the cells were analysed in a similar manner. In this case, the spectra were measured from the droplets dried on the surface of a quartz microscope slide using a dry objective 50× EC Epiplan-Neofluar LD, NA = 0.55 (Zeiss, Germany). Raman chemical maps were generated by multivariate decomposition of the cosmic-rays and baseline-corrected spectra into the spectra of the pure chemical components by using a *True Component Analysis* tool of the WITec Project Plus 6.1 software (Oxford Instruments - WITec, Germany).

### HPLC analysis

The extraction procedure involves the homogenization of *C. velia* cells using glass beads in a BeadBug™ microtube homogenizer, followed by the actual extraction. 0.1 M NaOH was used as the extraction solution (ES). 300 µL of ES was added to the extraction tube containing the dried and weighed sample. Three sizes (0.1-1.0 mm) of beads and 4 cycles per 10 seconds to the maximum speed were used to achieve complete homogenization. The homogenization tubes were then centrifugated at 12000 rpm per 10 minutes at 20°C using Hettich MIKRO 220. The collected supernatant was analysed by using HPLC FLD technique. 10 µL of the sample was injected through the column Kinetex F5 (2.6 µm pore size, 100 Å particle size, 150x4.6 mm) using Dionex UltiMate Autosampler RS. The temperature of the column was maintained at 35°C. Gradient chromatography was performed according to Gigliani et al. (2016) with slight modifications as follows: 0.1% (v/v) formic acid in water was used as mobile phase A (MPA) and methanol as mobile phase B (MPB). The flow rate was set to 0.6 ml/min. In the first 9 minutes, 100% MPA was eluted isocratically, then the proportion of MPB was linearly increased to 50% from 9 to 10 min and held until the 14^th^ minute. Subsequently, the MPB content decreased to 0% in the 15^th^ minute and was kept at zero until the 20^th^ minute to recondition the column. The Dionex UltiMate 3000 RS fluorescence detector was used to detect guanine at an excitation wavelength of 260 nm and an emission wavelength of 375 nm. Guanine was identified and quantified by comparison with a calibration curve generated from a purchased guanine standard (Sigma-Aldrich). In addition, the identity of the eluted compound was confirmed by spiking the samples with the standard.

### Scanning Electron Microscopy of whole cells

The first collection of *C. velia* cultures under nitrogen-repleted and -depleted conditions were collected for scanning electron microscopy by harvesting cells and centrifuged at ≈16000 g for 10 minutes. The fixation step was carried out in 2.5% glutaraldehyde in 0.1 M phosphate buffer. Later, the pellet was collected and post-fixed in 0.1% osmium tetroxide, followed by washing with F2 media. During the dehydration step, the pellet was treated with a graded series of acetone and dried with a critical point dryer. The samples were coated with gold and further observations were made using a JEOL JSM 7401-F scanning electron microscope.

### Transmission Electron Microscopy

Samples of *C. velia* were prepared for transmission electron microscopy by centrifugation at ≈16000 g for 10 minutes. The pellet was collected, washed with the F/2 medium, and subjected to a freeze substitution procedure. During freeze substitution, the samples were frozen in liquid nitrogen and then transferred to 2% OsO_4_ in anhydrous acetone, which served as a pre-cooled substitution medium and kept at -90°C for 72 hours. The samples were kept at -20°C and 4°C for 24 hours each. In the following, temperatures were increased by 4-5°C per hour until room temperature was reached. Samples were kept at room temperature and washed three times in pure anhydrous acetone and then embedded in Polybed 812 resin (with polymerization at 60°C for 48 hours). First, the samples were cut into 70nm ultra-thin sections with the help of an ultra-microtome using a diamond knife. To avoid the dropouts caused due to the ultra-thin sections, samples were cut into 90 nm thicker sections to retain the dropouts. The samples were then transferred to Formvar-coated grids and stained with uranyl acetate and lead citrate. Transmission electron microscopic observations were performed using a TEM (JOEL 1010) with an accelerating voltage of 80 kV.

### Isolation of guanine crystals

The isolation of guanine crystals was carried out by collecting 7-day old *C.velia* cells in a 50 mL of falcon tube. The sample was centrifuged using Hettich Mikro 120 centrifuge at 6000 rpm for 2 minutes and the supernatant was discarded. 1 mL of MilliQ water was added to the pellet and transferred to mortar and pestle. The sample was ground vigorously to disrupt the cells and transferred to the 1.5 mL Eppendorf tube. Centrifugation is carried at low-speed of 1500 rpm for 2 minutes. The supernatant was collected in a 1.5 mL Eppendorf tube and the pellet was discarded. The collected supernatant was again centrifuged at 8000 rpm for 2 minutes. The supernatant was discarded and the pellet is washed twice with MilliQ water by centrifugation at 8000 rpm for 2 minutes. To the pellet, 50 mL of MilliQ water was added and mixed thoroughly. 10 µL of the sample was pipetted on Superfrost^TM^ plus adhesion microscope slide and observed under polarized light microscope (Olympus DP 20 Model BX53, Shinjuku, Tokyo, Japan).

### Raman analysis of crystals isolated from C. velia cells after addition of a nitrogen source to the medium

The crystalline particles present in the cells of *C. velia* were extracted by density gradient centrifugation. Their presence in the centrifugation fractions was verified by polarized light microscope. To measure the Raman spectra, a drop (2 µL) was placed on a quartz glass slide where, after drying, it formed a ring at the edge of which crystalline particles were concentrated. These particles were Raman-mapped in the same way as the whole cells.

### Cryo-Scanning Electron Microscopy (cryo-SEM)

Cell pellets were centrifuged, transferred to 3 mm high-pressure freezing (HPF) carriers, which were filled with 20% w/v bovine serum albumin, and frozen using a Leica EM ICE (Leica Microsystems, Austria) high-pressure freezer. The samples were transferred under liquid nitrogen to a CryoALTO chamber (Gatan) pre-cooled to -160°C, fractured with a scalpel, freeze-etched at -98°C for 30 seconds and sputter-coated with gold. The samples were observed in a FESEM JEOL 7401F (JEOL Ltd.) at 1.5 kV with a working distance of 8 mm and a stage temperature of approximately -140 °C.

### Data analysis

The growth assessment of *C. velia* cells was conducted for each day from day 0 to day 7 under nitrogen-depleted and nitrogen-repleted conditions in triplicates. A graph is plotted using unpaired t-test, illustrating the correlation between optical density and the number of days using GraphPad Prism *v.*10. Crystals count was performed in approximately 30 cells. *C. velia* cells for each day from day 0 to day 5 under nitrogen-depleted and nitrogen-repleted conditions in triplicates. A box plot illustrating the average quantity of guanine crystals over time, is represented in the distribution of crystal numbers under nitrogen-depleted and repleted conditions. To evaluate variations in crystal counts under nitrogen-replete and nitrogen-deplete conditions throughout time points (Days 0–5), we initially conducted a normality assessment using the Shapiro–Wilk test for each day from day 0 to day 5 in nitrogen repleted and depleted conditions. On Day 1, Day 3, and Day 4, the data exhibited a normal distribution (p > 0.05), prompting the use of an unpaired t-test to compare group means. Conversely, for Day 0, Day 2, and Day 5, the data failed to satisfy the normality criteria (Shapiro–Wilk p ≤ 0.05), prompting the use of the Wilcoxon rank-sum test (Mann–Whitney U test) for comparing group medians. All statistical analyses were conducted using R software *v*.4.4.2, with a significance level established at p < 0.05.

### Raman spectroscopy, electron microscopy and EDX analysis

Standard guanine powder (98% A12024, Thermo scientific) and guanine crystals isolated from *C. velia* were analyzed by Raman spectroscopy using an upright Raman microscope (WITec Alpha 300, Oxford Instruments) with 532 nm laser excitation. Raman spectra were acquired using a UHTS 300S Green–NIR spectrometer equipped with a 600 gr/mm grating. The same samples were mounted on SEM stubs and imaged without conductive coating using a VERIOS 5 UC scanning electron microscope (Thermo Fisher Scientific). Imaging was performed at an accelerating voltage of 3 kV for topographic acquisition and 5 kV for energy-dispersive X-ray (EDX) analysis using a Quantax FlatQUAD detector (Bruker), with a probe current of 13 pA.

A pelleted suspension of *C. velia* was high-pressure frozen using a EM ICE system (Leica Microsystems). Frozen samples were transferred to a Quorum cryo-sample preparation system, where cells were freeze-fractured at −140 °C and subsequently freeze-sublimated for 2–5 min at −95 °C. Samples were then imaged at -1401C using the same SEM at an accelerating voltage of 1 kV and a probe current of 13 pA. EDX analysis was performed at an accelerating voltage of 2 kV.

## Results and Discussion

### Physiological role of crystalline guanine

Cells cultivated in F/2 media for 7 days were transferred to nitrogen-repleted and nitrogen-depleted conditions and allowed to grow for one week to observe cell density (Figure 1). Under nitrogen-depleted conditions, no significant difference was observed from day 0 to day 2 and from 3 to day 7, significant differences were observed (p <0.05). These results confirm that the growth rates were restricted under the nitrogen-depleted condition compared to nitrogen-repleted conditions (Figure 2b).

**Figure 2:**
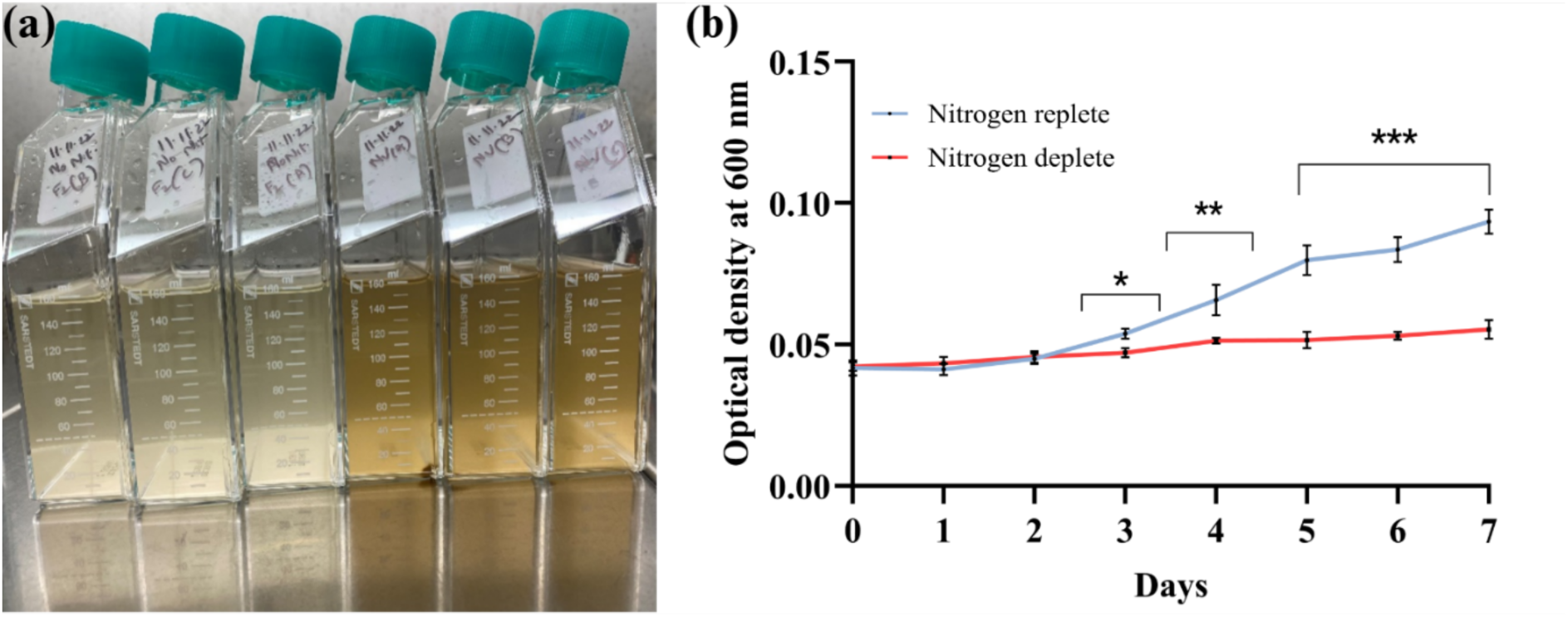
Growth of *Chromera velia* cells in response to nitrogen availability. a) *Chromera velia* cultures after 7 days of cultivation in nitrogen deplete (left three vials) and nitrogen replete (right three vials) F/2 medium. b) Growth curves of *C velia* cultivated in nitrogen replete and deplete conditions for 7 days. Optical densities were plotted as average and standard deviation of triplicate cultures in both conditions. No significance difference in growth between nitrogen repleted and depleted conditions was observed from day 0 to day 2 (p >0.05), while from day 3 to day 7 significant differences were observed (p <0.05). Brackets with asterisks indicate the p values for the specific days with (*p<0.05, **p<0.01, ***p<0.001).

Cells from the cultures were subjected to polarised light microscope. The guanine crystals remained unchanged under nitrogen repleted conditions (Figure 3a). However, under nitrogen-depleted conditions, a gradual decrease in crystal presence was observed from day 1 to day 4, and by day 5, crystals had completely disappeared in the *C. velia* cells (Figure 3b). When these cells were transferred back to culture media with nitrogen-repleted conditions, the reappearance of guanine crystals was observed within 24 hours (Figure 3c). The results from the Raman spectra maps of *C. velia* cells under nitrogen repleted conditions confirmed that the crystals are composed of anhydrous guanine in beta-polytype. Lipido-proteins and a polyene were also identified under nitrogen-repleted conditions (Figure 3d). However, under nitrogen-depleted conditions, starch was identified instead of guanine, together with neutral lipids and polyene (Figure 3e). A total of 30 cells were counted in triplicate for both conditions from day 0 to day 5 to determine the average number of crystals in *C. velia* cells. A boxplot comparing nitrogen-repleted and -depleted conditions with the number of days (x-axis) and the average number of crystals in cells (y-axis) shows the dependence of the number of crystals on the availability of nitrogen in the medium, or on the growth time in nitrogen-free medium. The number of crystals under nitrogen-repleted conditions remains similar from day 0 to day 5 (∼ 6 crystals per cell), while under nitrogen-depleted condition, the number of crystals decreases from day 2 to day 5 and guanine crystals were completely absent on day 5 (Figure 3f). We quantified the guanine content from the extracts of *C. velia* cells in nitrogen depleted and repleted conditions using HPLC. The guanine content is higher in nitrogen replete condition compared to the nitrogen depleted condition (Figure 3g).

**Figure 3:**
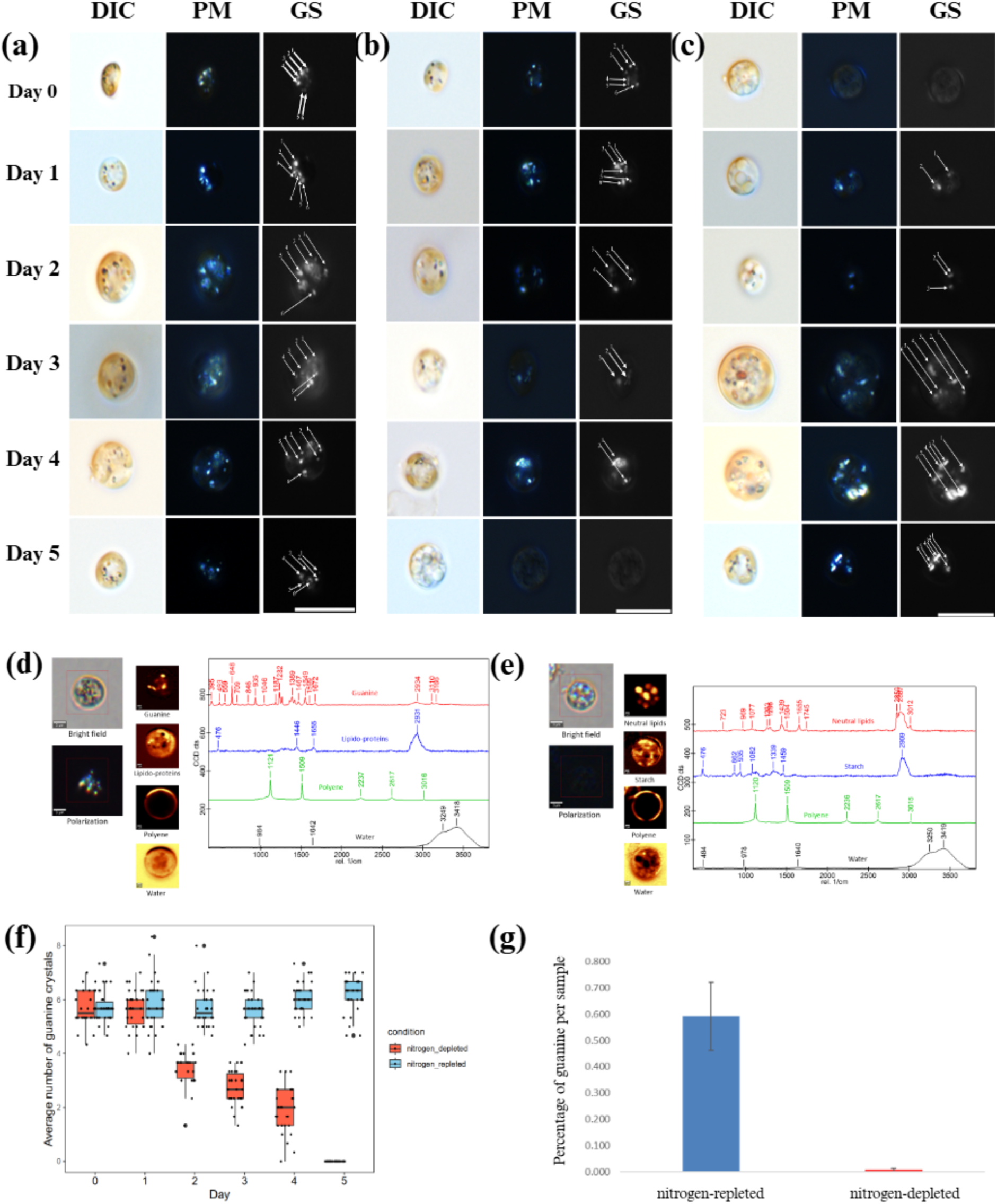
Guanine crystal dynamics in *Chromera velia* in response to nitrogen availability. a) Cells grown in nitrogen replete medium; Differential interference contrast (DIC) and polarised light (PM) microscopy images; guanine crystals are observed from day 0 to day 5 (compare to Figure 1). Arrows on the grayscale (GS) images mark individual crystals as they were counted. b) Cells kept in nitrogen-deplete medium from day 0 to day 5. Decrease in numbers of guanine crystals was observed from day 3, and on day 5, the total absence of crystals was noted. c) Reappearance of guanine crystals upon transfer of the cells to nitrogen replete medium was observed from day 1. d) Brightfield and polarisation micrographs, along with Raman chemical maps revealed the distribution of crystalline guanine, lipido-protein mass, an unknown polyene located in the cell wall, and water in a typical *C. velia* cell after 5 days of cultivation under nitrogen replete conditions Scale bar = 3 µm. e) Brightfield, polarisation micrographs and Raman chemical maps in nitrogen depleted conditions of showing the distribution of neutral lipids, starch grains, an unknown polyene located in the cell wall, and water in a typical *C. velia* cell cultured under nitrogen-depleted conditions for 5 days. Scale bar = 3 µm. f) Box plot showing average guanine crystals counts from day 0 to day 5 under nitrogen repleted and depleted conditions. X-axis represents days, and Y axis represents the number of guanine crystals per cell (n = 30 cells per condition and day) in *C. velia* cells under nitrogen repleted and depleted conditions. Color legend: Blue boxes represent nitrogen repleted and red boxes represented nitrogen depleted conditions. Horizontal lines represent the mean; whiskers indicate the variability outside of the upper and lower quartiles, and black dots represent outliners. g) Guanine content as measured by HPLC in *C.velia* cells under nitrogen repleted and depleted conditions, mean and standard deviations of the triplicate samples in both conditions are shown.

In other organisms, nitrogen starvation not only has been shown to promote the degradation of the guanine crystals (in the marine dinoflagellate *Amphidinium carterae* (Mojzeš et al., 2020) and in the green alga *Chlamydomonas reinhardtii* (Goodeneoughet al. 2025),but also lead to a decrease in biomass, with an increase in the formation of starch granules and lipids droplets (in the unicellular green alga, *Dunaliella tertiolecta,* the diatom, *Phaeodactylum tricornutum,* and the eustigmatophyte *Nannochloropsis oculata* (Şirin & Serdar, 2024)). In green microalgae *Neochloris oleoabundans*, starch and lipids accumulated in both salt and freshwater during nitrogen starvation (Jaeger et al., 2018). In diatoms such as *Phaeodactylum tricornutum,* nitrogen stress conditions lead to an increase in monosaturated palmitoleic acid and the accumulation of lipids (Liu et al., 2020). Studies shown that *Chlamydomonas reinhardtii* shows an increase in the starch content upon nitrogen starvation (Siaut et al., 2011) and the accumulated starch was converted into lipids (Wase et al., 2014). A study led by Findinier et al. (2019) demonstrates the importance of *BSG1* gene in starch metabolism that led to switch from pyrenoidal to storage starch mechanism. The deletion of *BSG1* gene in *Chlamydomonas reinhardtii* resulted in reduction rate of starch degradation and modified structure of amylopectin when observed under electron microscopy. When starch granules are extracted, the diameter of each starch granules were determined from 0.5 to 5 µm using scanning electron microscopy. Under nitrogen starvation conditions, they accumulated abnormally two different sizes, small and large starch granules, irregular in shape and twice in size when compared to wild-type granules which has only one type of granules of regular size distribution (Findinier et al., 2019). In *C. velia*, absence of nitrogen led to suppression of growth from day 3 to day 7 (Figure 2b). In addition, Raman microspectroscopy of *C. velia* cells under nitrogen depleted conditions, revealed an increase in starch and neutral lipids (Figure 3e), confirming that the absence of nitrogen promotes the formation of starch and lipids, as previously mentioned for other algae.

The surface morphology of *C. velia* cells was examined by SEM under nitrogen-repleted and -depleted conditions (Figure S4). The cell size under both conditions showed no variation throughout the experiment. The surface morphology of *C. velia* cells in nitrogen-repleted media showed no significant changes (Figure S4).

Transmission electron microscopy (TEM) was used to localise the crystals in cells of *C. velia*. The TEM micrographs showed the distribution of vacuoles or storage granules in the cytoplasm of *C. velia* under both nitrogen-repleted (Figure S5 a, b) and depleted conditions (Figure S5 c, d). The samples were prepared by ultra-thin sections (70 nm), which resulted in frequent dropouts or holes in the *C. velia* cells under both nitrogen-repleted and depleted conditions (Figure S5). Similar observations of holes instead of crystal inclusions or guanine storage vacuoles have been made with ultra-thin sections of *Chlamydomonas reinhardtii* (Goodenough et al. 2025). To retain the holes or dropouts in the cells, thicker sections were prepared (90 nm) and the samples were transferred to formvar-coated grids and stained with uranyl acetate and lead citrate. When the samples were observed under TEM, *C. velia* cells under nitrogen depleted conditions revealed the presence of vacuoles or storage granules distributed throughout the cytoplasm. The vacuoles are surrounded by the large lipid droplets and plastids (Figure 4 a, b). On the other hand, under nitrogen repleted conditions, *C. velia* cells shows a lesser number of vacuoles or storage granules in comparison to nitrogen depleted conditions along with clearly visible nucleus and pyrenoid present in the plastid (Figure 4 c, d).

**Figure 4:**
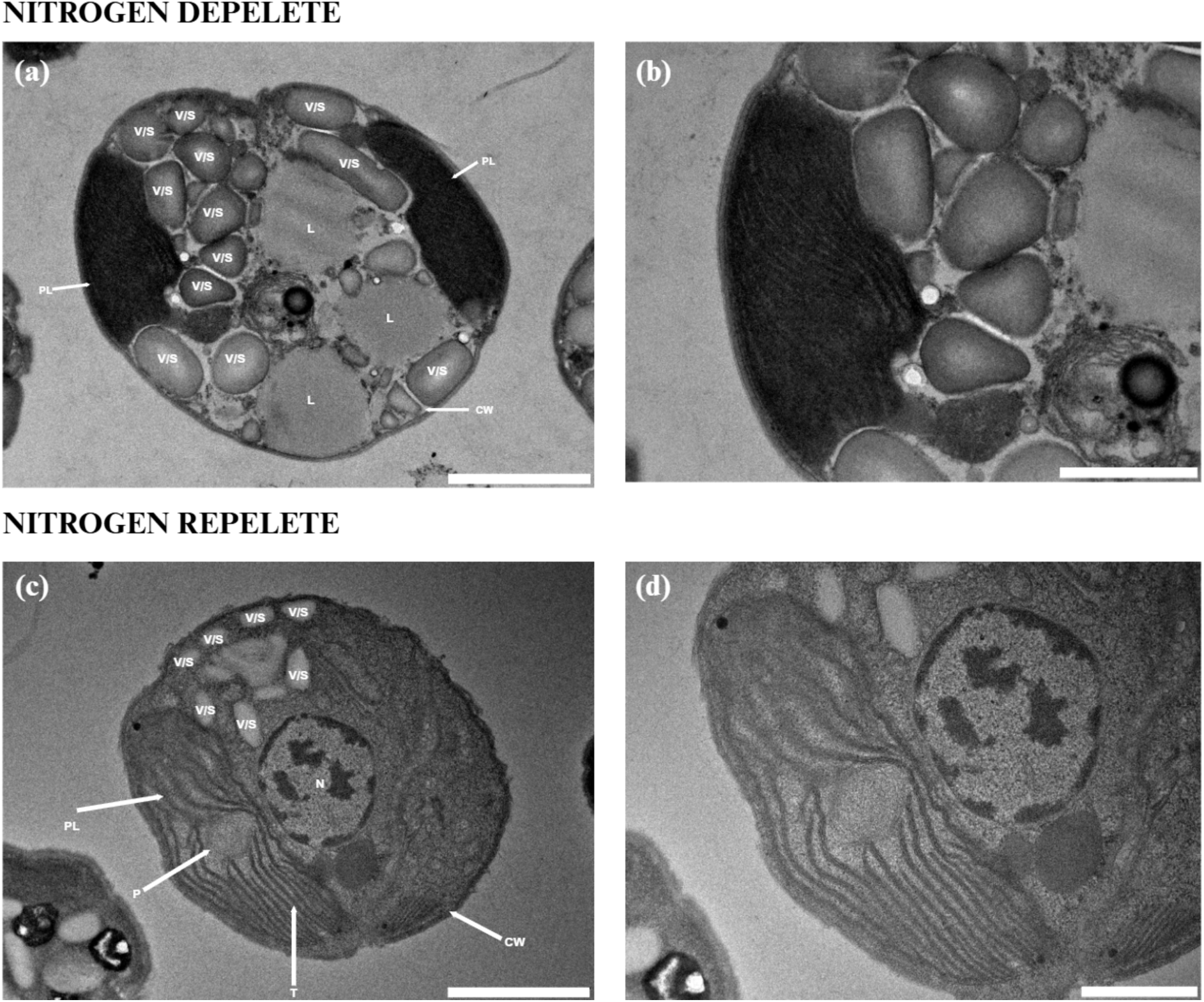
Transmission electron micrographs of *Chromera velia* cells grown in nitrogen deplete and replete conditions. Images were taken from thicker sections of 90 nm, a-b) nitrogen deplete conditions, c-d) nitrogen replete conditions b) and d) are enlargements of a) and c) respectively. CW: cell wall, PL: plastid and L: lipids, V/S: vacuoles or storage granules. Scale bars: a) 2 μm, b) 500 nm, c) 2 μm and d) 500 nm.

The morphological and optical characteristics of guanine crystals and starch granules differ when observed under a polarised light microscope. Guanine crystals are smaller, plated and brightly coloured (Wagner et al., 2023; Pinsk et al., 2022). In contrast, starch granules are semi crystalline in various sizes from 0.1 to 100 µm in diameter and appears in ovoid, ellipsoidal, spherical, angular in shapes (Findinier et al., 2019), exhibiting unique birefringence patterns called the Maltese cross determined under polarised light microscopy. The Maltese cross is the hallmark of starch granules, which appears as dark pattern, divided into four light quadrants (Evans et al. (2003) The cross formation is due to the hilum, a central point or core that serves as a starting point for the development of starch granule (Miranda Joyce Aparecida Tavares de et al., 2019). However, the Maltese cross does not appear when the guanine crystals were observed under the polarised light microscopy showing clear differentiation among the guanine crystals and starch granules crystals.

### Crystals in zoospores of Chromera velia

Zoospores from a 7-day-old culture of *C. velia* were isolated according to the method described in Richtová et al. (2023) and fixed in 4% paraformaldehyde. The isolated zoospores were observed using fluorescence microscopy, differential interference contrast image, plastid autofluorescence image were taken, along with polarised filters. Birefringent structures could be observed under polarised light illumination (Figure 5 h, k, n). The isolated zoospores from *C. velia* cells were subjected to Raman micro-spectroscopic analysis confirming the presence of guanine crystals (Figure 5p, q).

**Figure 5:**
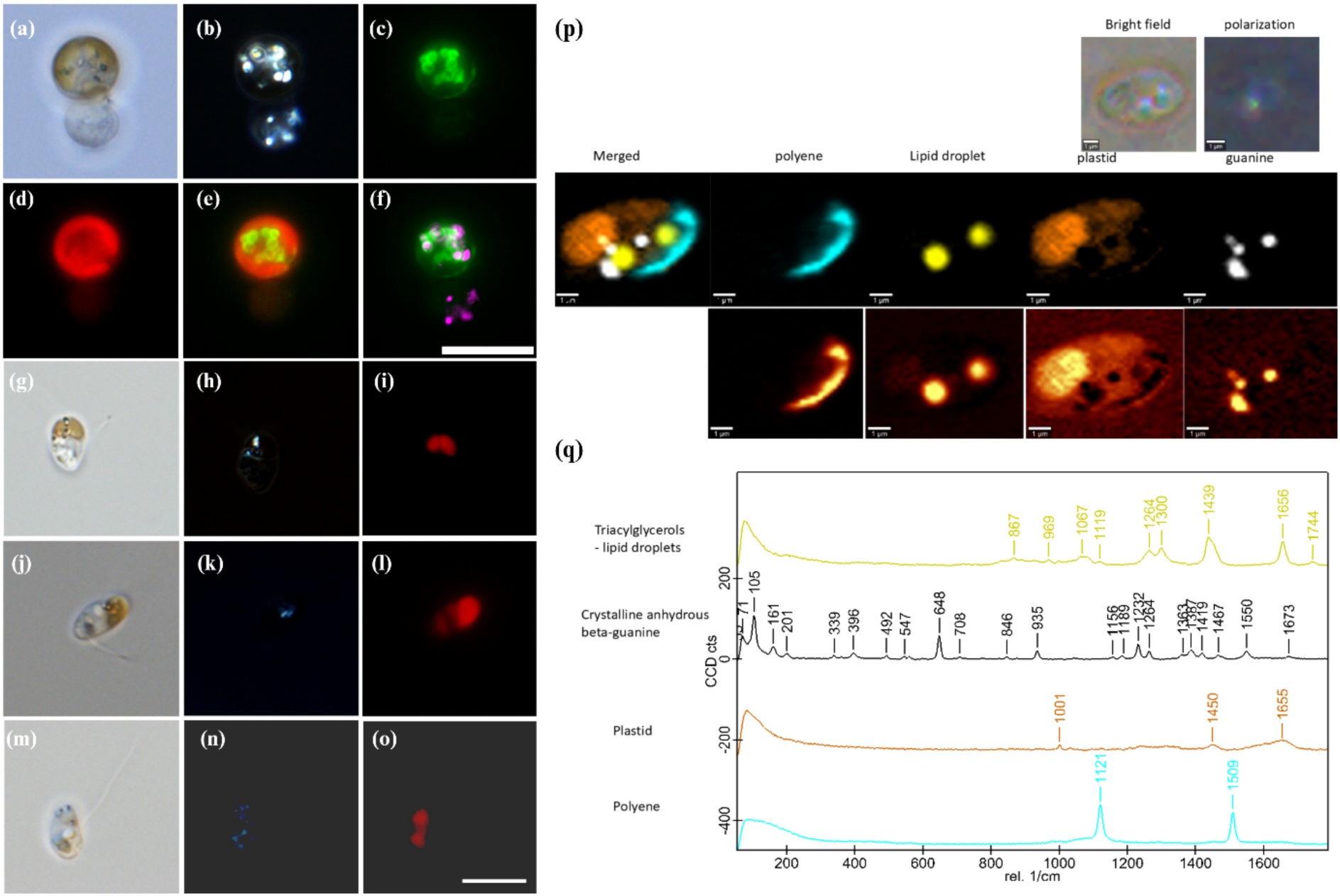
Vacuolar staining, visualisation of guanine crystals and identification of guanine crystals in zoospores of *Chromera velia.* a) Bright-field microscopy image of *C. velia* cells stained with Lysotracker ^TM^ Green DND-26, which accumulates in acidic vacuolar compartments. b) Polarized light micrograph highlighting the birefringence of guanine crystals, which appear as bright, refractile structures. c) Fluorescence microscopic image revealing the specific labelling of the vacuoles stained by DND-26. d) Autofluorescence of the plastids. e) Merged image of c) fluorescent staining and d) plastid autofluorescence, revealing the spatial distinction between guanine containing vacuoles and plastids. f) Merged image overlaying b) polarised light birefringence of guanine crystals and c) DND-26 staining of the vacuoles (magenta) indicating localisation of birefringent guanine crystals within the labelled vacuoles, Scale bar = 10 μm. g-o) *Chromera velia* zoospores isolated by the method described in Richtova et al; 2023 and fixed in 4% paraformaldehyde. Images g, j, m) Zoospores observed under differential interference contrast. Images h, k, n) observed under polarized light microscopy highlighting the birefringence of guanine crystals and images (i, l, o) shows the autofluorescence of plastids in the same cells. Scale bar =5 µm. p) Isolated zoospores from Chromera velia cells under bright field, polarised and Raman microspectroscopy. q) Raman spectrum maps depicting the presence of lipid droplets, crystalline anhydrous guanine, plastid and polyene.

### Intracellular arrangement of guanine crystals in Chromera velia

The observed guanine crystals can be seen moving within the cytosol of *C. velia* in polarizing light microscopy (Supplementary data S1), indicating that individual guanine crystals, which on their own would be below the resolution limit for light microscopy, are ordered to coordinated aggregates, which are following the same movements per aggregate, however, individual aggregates are moving independent from each other in the cell. We therefore set out to clarify the cellular arrangement of the crystals, and in freeze etching and freeze fracture techniques, were able to confirm the location of the crystalline material within small vacuoles (Figure 8, S2). The vacuoles were approximately 0.5-1 µm in size, while the individual crystals were approximately 0.2 µm long (Figure S3).These crystals were furthermore shown to be nitrogen enriched compared to their surrounding via EDX elemental analysis (Figure 6).

**Figure 6:**
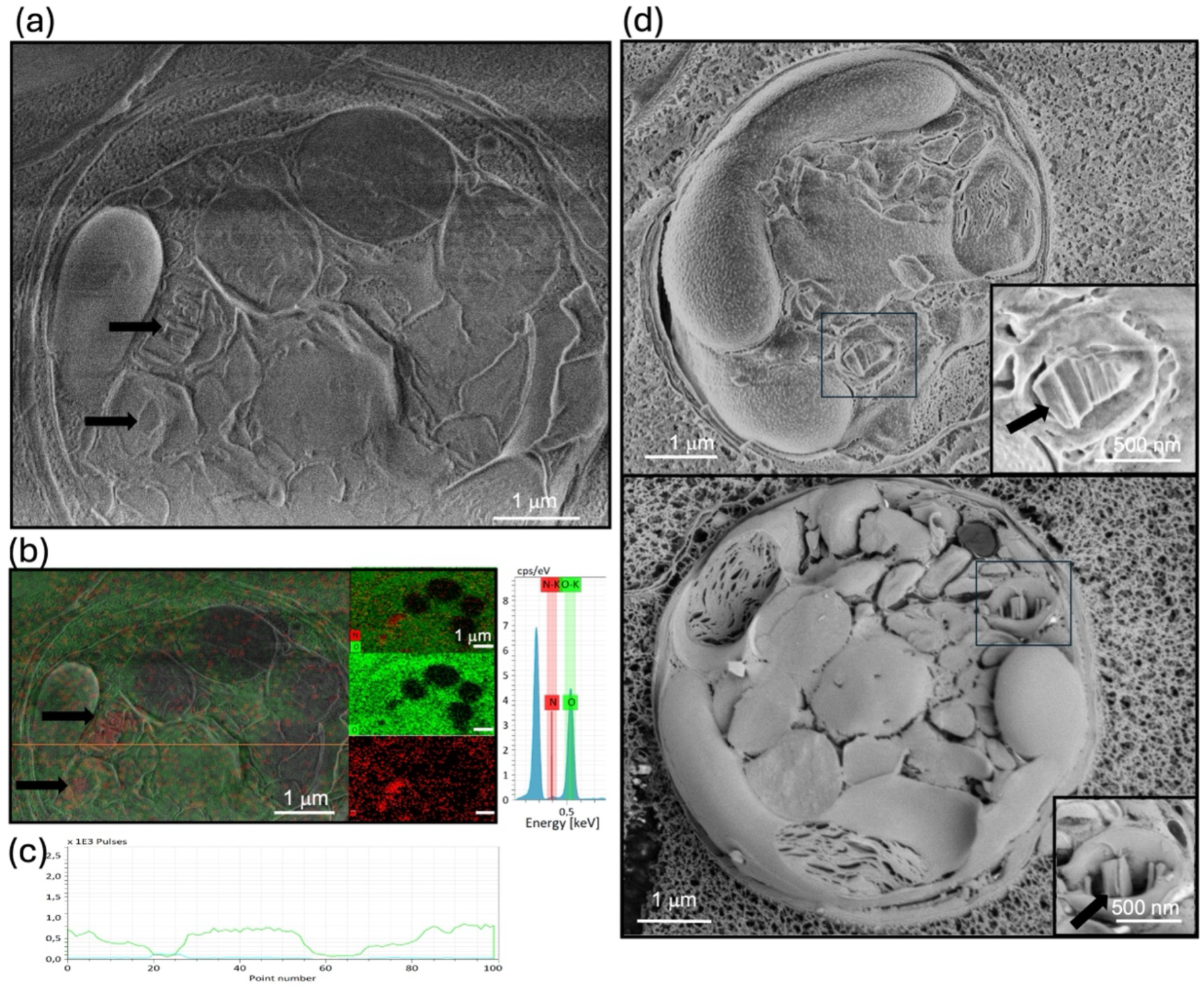
Cryo-SEM–EDS analysis of *Chromera velia* cells showing intracellular guanine crystals (black arrows, details in insets). Topographic images (a, d) were acquired on a Verios 5 UC at 1 kV, 13 pA. EDS elemental maps and the corresponding spectrum (b), confirming the crystal composition of the structures shown in (a), were acquired at 2 kV using a Quantax FlatQUAD EDS detector. The distribution of N along the orange line in (b) is shown in graph (c).

For independent confirmation, we stained cells with the green dye Lysotracker DND-26, an acidotropic probe consisting of a fluorophore linked to a weak base that is partially protonated at neutral pH. Upon diffusion into acidic compartments, the weakly basic moiety is protonated and subsequently binds to the membrane, thus making the structure of the stained organelle visibly distinct. We observed that upon excitation, the membrane-bound fluorophore emitted fluorescence that appeared as spherical structures surrounded by the plastid of *C. velia* cells (Figure 5 a-f). The localization of the identified structures shows a strong correlation with the position of guanine crystals. However, it should be noted that this colocalization may not be entirely precise and clear in Figure 5 as the guanine crystals were mobile within the cells and the staining method employed was only compatible with live cells, making it impossible to fix the samples for more stable imaging. Cells treated solely with either Triton-X100 or DMSO (with no Lysotracker staining) showed no fluorescence when using a GFP filter. These results are in line with the previous studies by Hirsch et al. (2015) and Jantschke et al. (2019) showing that beta-polytype anhydrous guanine crystals are biogenic forms of guanine that are vertically stacked in irregular shapes by hydrogen bonding (Hirsch et al., 2015; Jantschke et al., 2019). In a comprehensive study by Pilátová et al. (2022), more than 200 microbial species from different eukaryotic groups were examined for the presence of crystalline inclusions using polarised light microscope and Raman Analysis. They proved that anhydrous guanine is the most dominant form of storage guanine. Moudříková, et al. (2017) found beta-polytype guanine crystals in the microalgae *Desmodesmus quadricauda* and *Trachydiscus minutus* are spherical in shape and compactly arranged in the vacuoles. In dinoflagellate *Calciodinellum. operosum aP.,* vacuoles range from 0.5 to several micrometres, contain crystals in the size range of 100 - 800 nm, which were identified on the inner sides of the chloroplast, suggesting light utilisation or UV protection using cryo-SEM and FIB-SEM. Using Raman microspectroscopic analysis, crystals inside the vacuoles are identified as anhydrous (Jantschke et al., 2019). Research led by Shebanova et al. (2017), identified two different types of vacuolar inclusions, in *Chlorella vulgaris*, *Parachlorella kessleri* and *Desmodesmus* sp. each with specific functions using EDX analysis. Type I inclusions store phosphorus in the form of polyP, while type II is involved in nitrogen storage, resulting from the acclimatisation of nitrogen starvation and high CO_2_ levels (Shebanova et al., 2017). The formation of guanine crystals was also observed by Mojzeš et al., (2020) in the marine dinoflagellate *Amphidinium carterae* that showed, similar to *C. velia*, an accumulation of guanine inclusions in the nitrogen rich media while during nitrogen starvation, guanine was degraded to supply nitrogen for the essential metabolic processes.

### Isolation of guanine crystals from C. velia

Isolation of crystals from *C. velia* cells cultivated in nitrogen repleted conditions was achieved using a newly developed protocol (Figure 7). The isolated crystals when subjected to polarized light microscopy, where we observed the birefringence of the crystals (Figure 7a-d). The isolated crystals furthermore contained a similar level of nitrogen enrichment as the crystalline inclusions in freeze fracture images of whole cells (compare Figures 6 and 9), and Raman micro-spectroscopy showed that they consist of anhydrous guanine crystals in beta-polytype (Figure 10).

**Figure 7:**
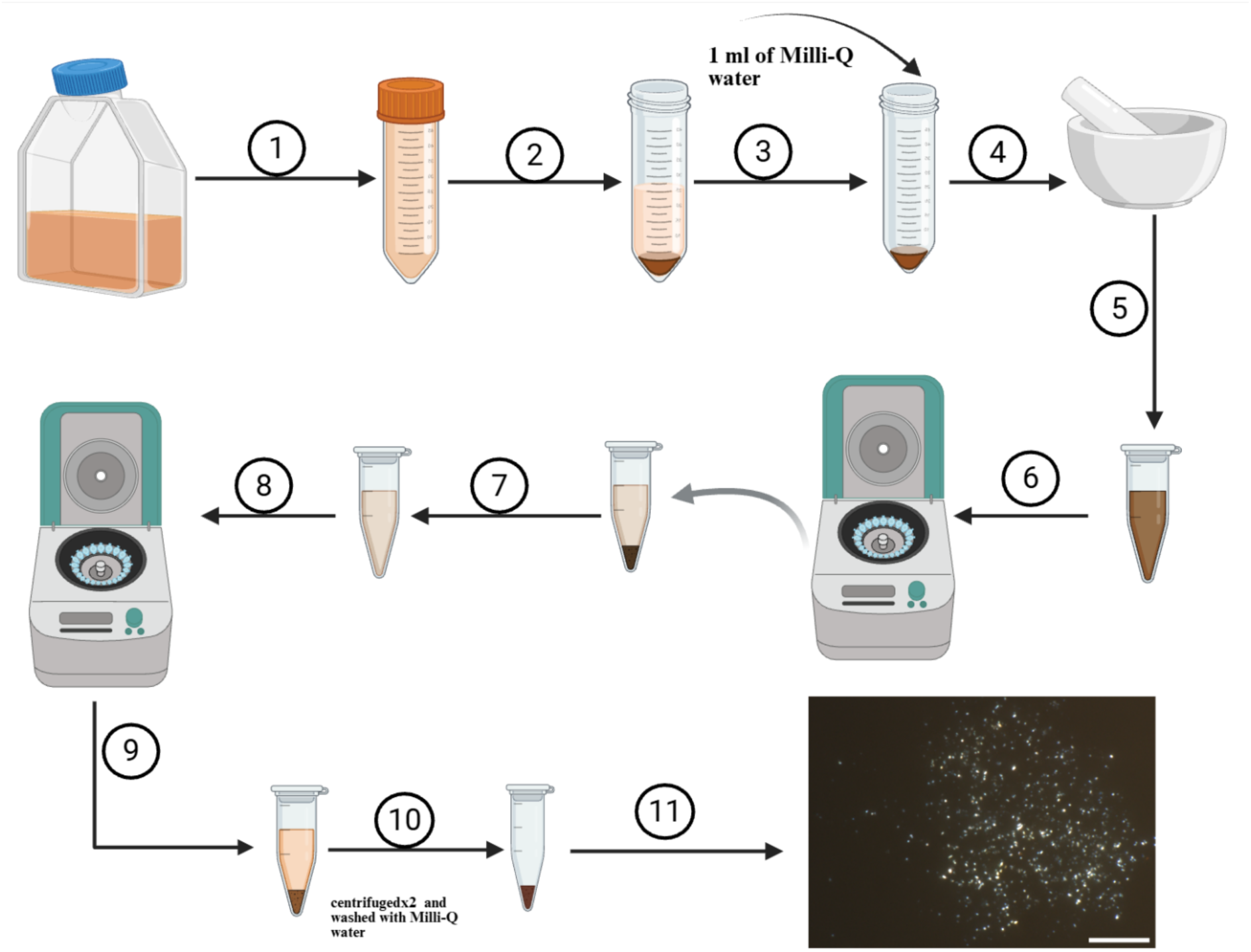
Isolation of guanine crystals from *Chromera velia* cells. 1) *Chromera velia* cells were transferred to 50 ml falcon tube. 2) cells were subjected to centrifugation at 2000 × g for 2 minutes. 3) Supernatant was discarded and 1 ml of Milli-Q water was added to the pellet. 4) The sample was collected and transferred to mortar and pestle. 5) after vigorous grinding, the sample was collected in centrifuge tube and 6) subjected to centrifugation in low speed at ≈180 × g for 2 minutes. 7) Supernatant was collected and the pellet was discarded. 8) The collected supernatant is subjected to centrifugation at ≈5000 × g for 2 minutes. 9) After centrifugation, the sample was separated as pellet and supernatant. 10) Supernatant was discarded and the pellet was washed with Milli-Q water by centrifugating the sample at ≈5000 × g for 2 minutes twice. 11) 10 ul from the pellet was added on the microslide and guanine crystals were identified under polarised light microscopy. Scale bar is 20 µm.

**Figure 8:**
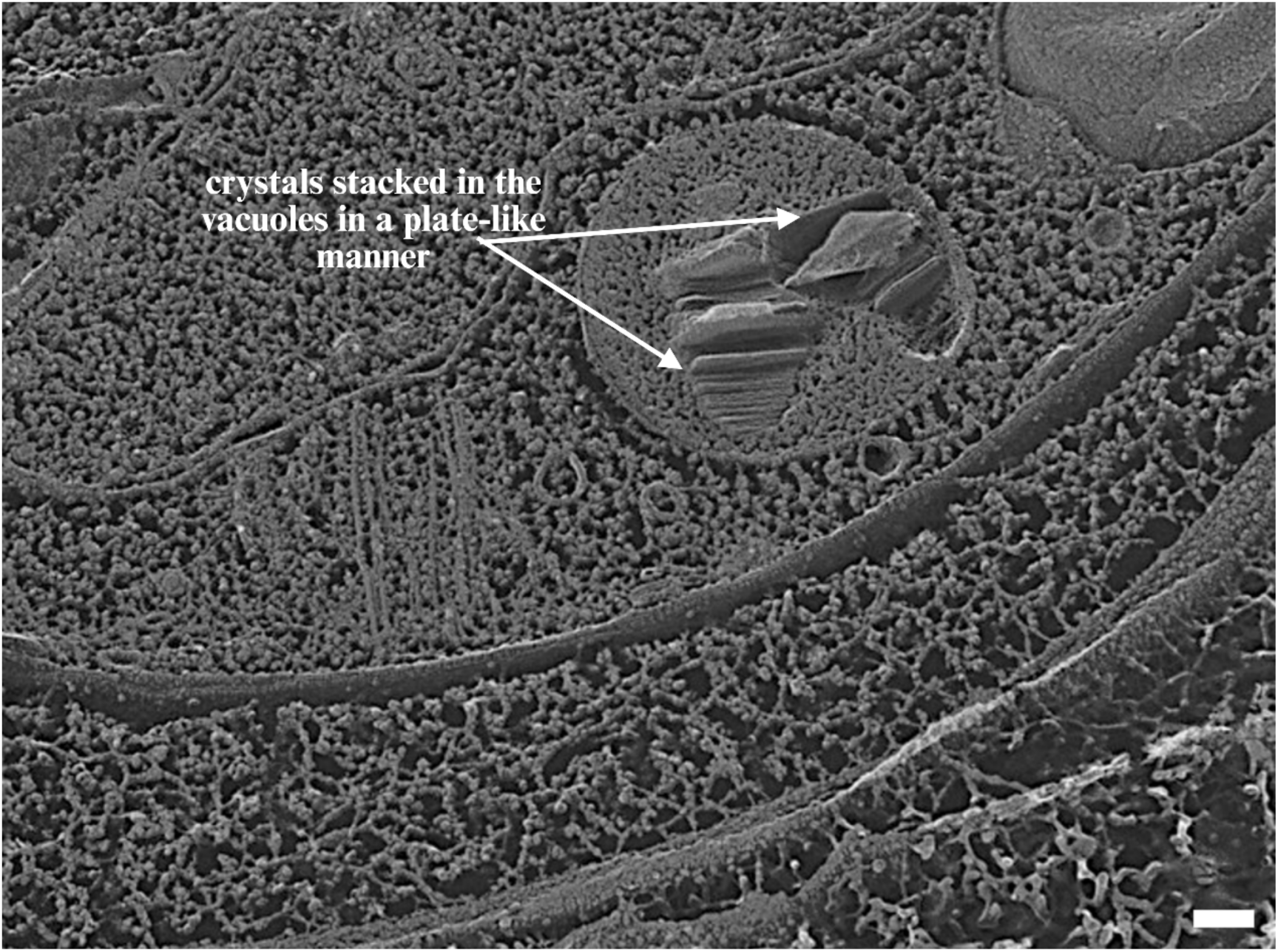
Freeze etching image of *Chromera velia*. Stacks of plate-shaped crystals are observed within vacuoles in the cytosol of *Chromera velia*, scale bar = 100 nm.

**Figure 9:**
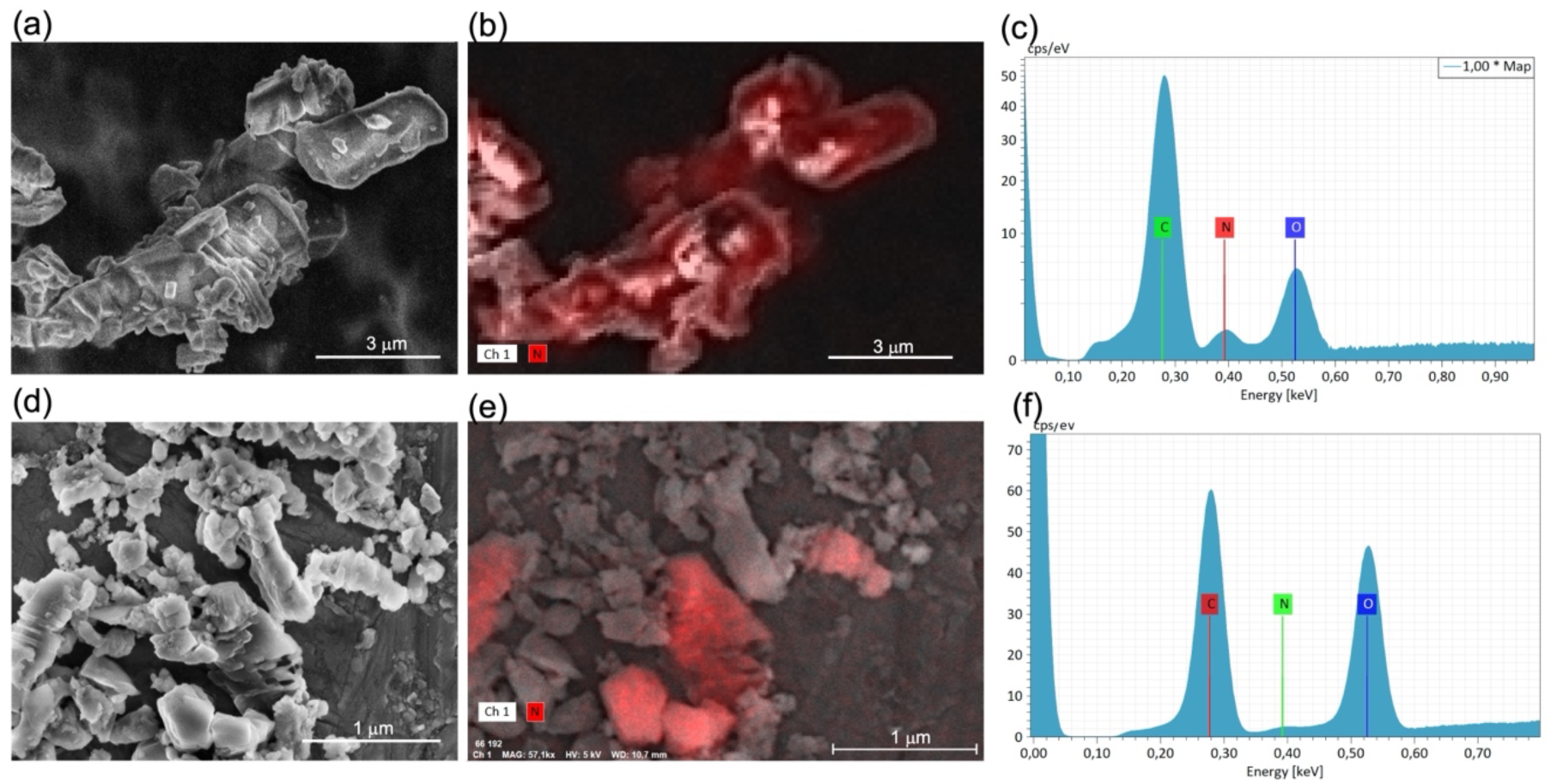
Cryo-SEM-EDS of commercial guanine powder standard (a-c) and guanine crystals isolated from *Chromera velia* cells (d-f). Topographical images were acquired using Verios 5UC at 3 kV(A, D). EDS elemental maps (b, e) and spektra (c, f) were acquired at 5 kV using Quantax FlatQUAD detector.

**Figure 10:**
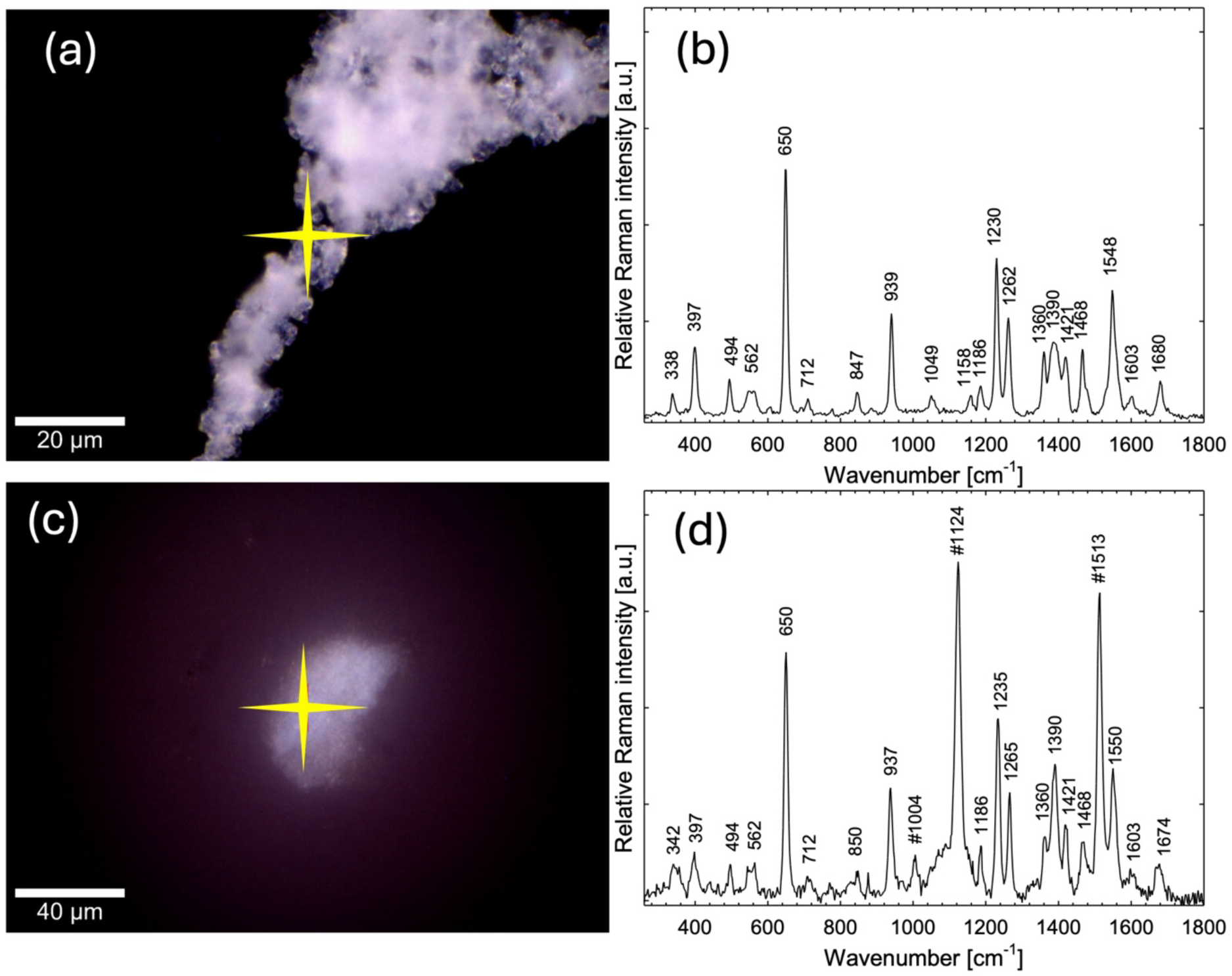
Bright-field images and Raman spectra of reference guanine crystals (a, b) and guanine crystals isolated from *Chromera velia* cells (c, d). Raman spectra (b, d) were acquired using a 532 nm excitation laser at 50 mW. Yellow crosses in (a) and (c) indicate the positions from which the single-point spectra shown in (b) and (d) were collected. Extra Raman bands at 1124 and 1513 cm^-1^, observed only in the spectra of *C. velia* guanine crystals, are due to an unidentified polyenic compound present in the cell walls. The band at 1004 cm^-1^ corresponds to phenylalanine vibrations of proteins.

## Conclusion

Taken together, our results suggest that the crystalline guanine found in the alveolate alga *Chromera velia*, is located within specialized guanine storage vacuoles, which play an important role in the storage and metabolism of nitrogen in the cell. Guanine (or other purine) crystals have recently been identified in a high diversity of microorganisms including algae with primary plastids (e.g. *Chlamydomonas reinhardtii*) or with complex plastids (e.g. in members of the *Stramenopiles*). The similarity between the guanine storage vacuoles, despite deep phylogenetic branches between the organisms in which they are found, might indicate that the integration of solid-phase guanine in the cellular nitrogen metabolism evolved early in the radiation of eukaryotes.

## Acknowledgements

This work was supported by the Czech Science Foundation (21-26115S, VGG, JP, DE, PM, AG and 23-06203S, MO) and by the Institute of Parasitology of the Czech Academy of Sciences. Part of this work was carried out with the support of ELIXIR CZ Research Infrastructure (ID LM2023055, MEYS CR, AG). The work of JP at the Molecular Foundry was supported by the Laboratory Directed Research and Development program at Lawrence Berkeley National Laboratory (LDRD 25-116), the Office of Science, Office of Basic Energy Sciences, of the U.S. Department of Energy under Contract No. DE-AC02-05CH11231. The work of UG was supported by grants DE-SC0006873 and DE-EE0003046 from the US Department of Energy and grants from the International Center for Energy, Environment and Sustainability, Washington University. We acknowledge the BC CAS core facility LEM supported by the Czech-BioImaging large RI project (LM2023050 and OP VVV CZ.02.1.01/0.0/0.0/18_046/0016045).

## Supplementary Material

**Movie S1:**
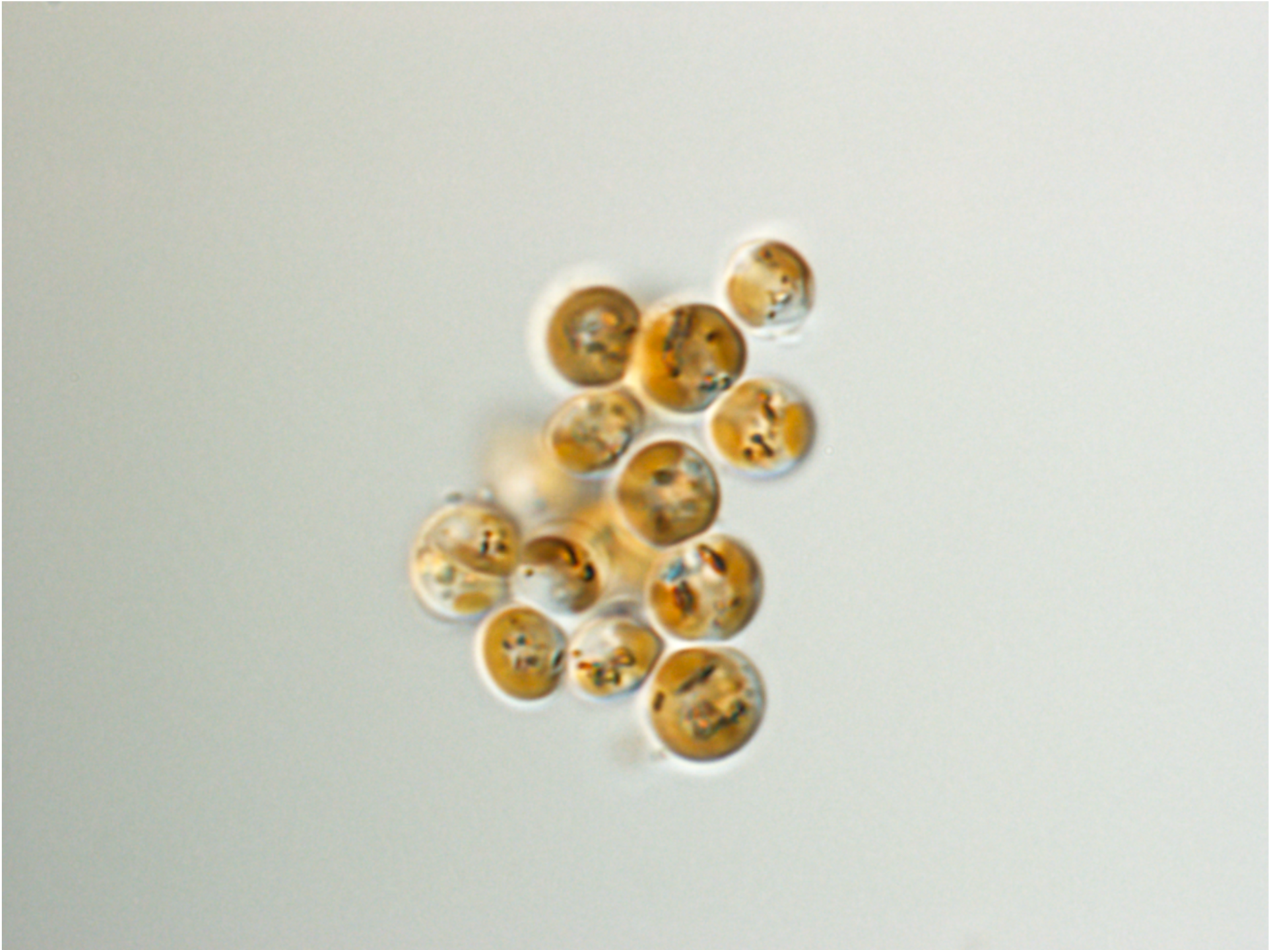
In vivo imaging of bifringent crystals in *Chromera velia*. The movie is shown in real-time, illumination is blended from DIC (beginningof clip) to polarized light imaging (end of clip) by rotation of the DIC prism. (see file “Gonepoguetal_S1.avi”, screenshot shown above).

**Figure S2:**
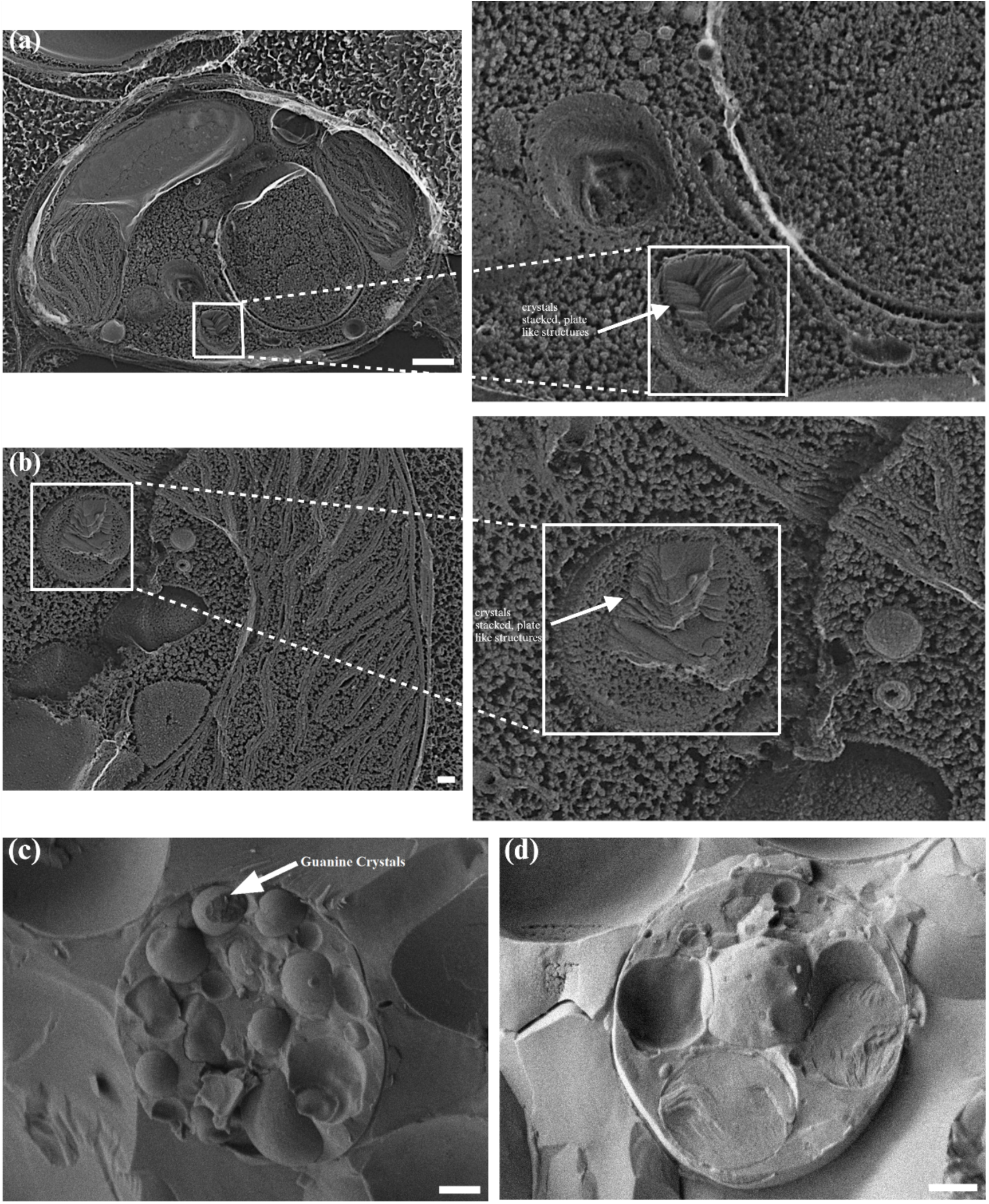
Freeze etching and Freeze fracture micrographs of Chromera velia cells showing the presence of crystals arranged in a stack of plate like manner inside the vacuoles. Image (a) and (b) are freeze etched, (c) and (d) are freeze fractured without etching, scale bars (a) = 500 nm, (b) = 100 nm, (c, d) = 1 µm.

**Figure S3:**
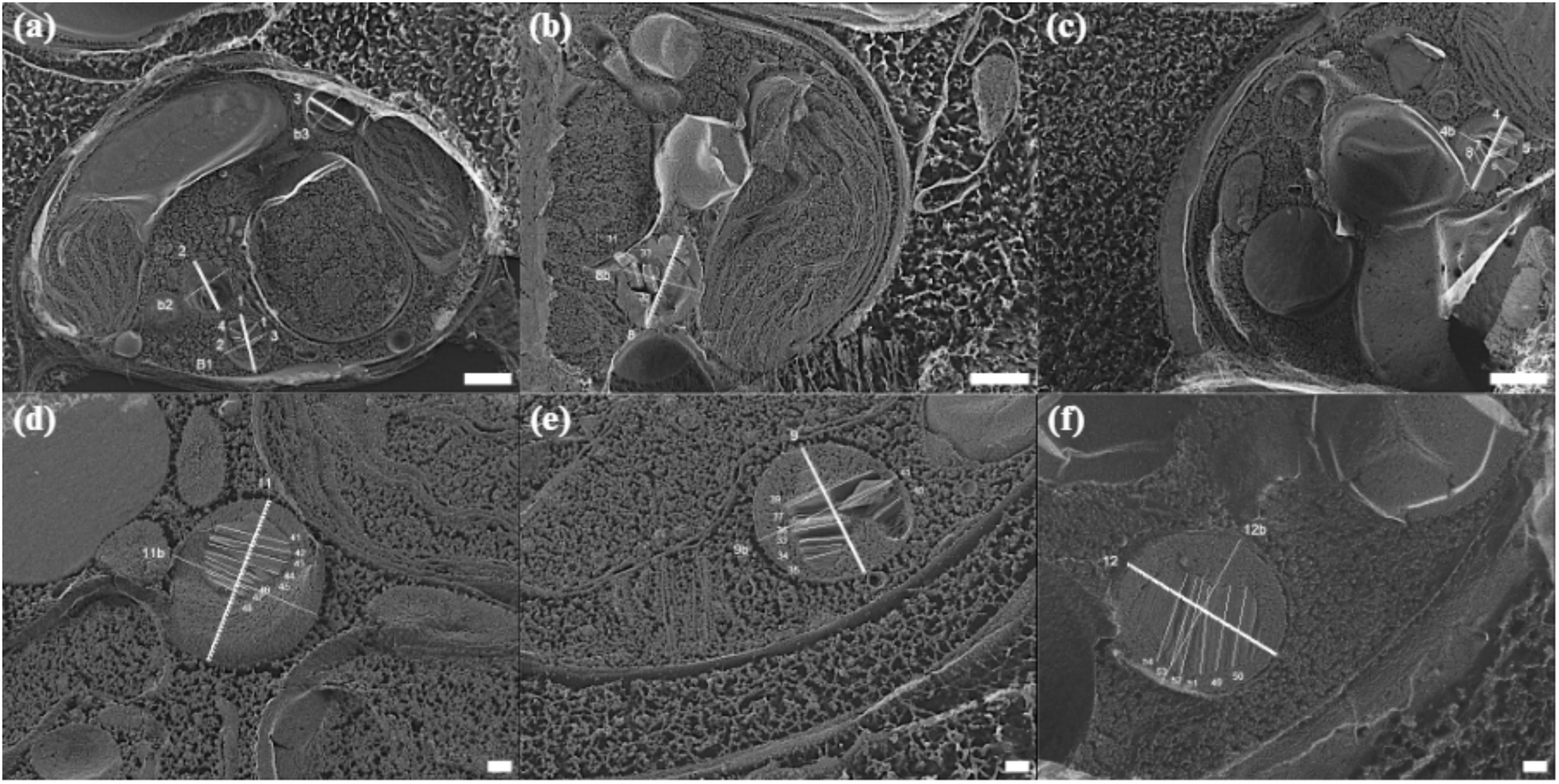
Size measurements on freeze etching micrographs of *Chromera velia* cells. Vacuole lengths and widths are marked with lines in images a-f), each crystal inside the vacuole is labelled with lines and numbers in images a-f). Scale bars a-c) = 500 nm and d-f) = 100 nm.

**Figure S4:**
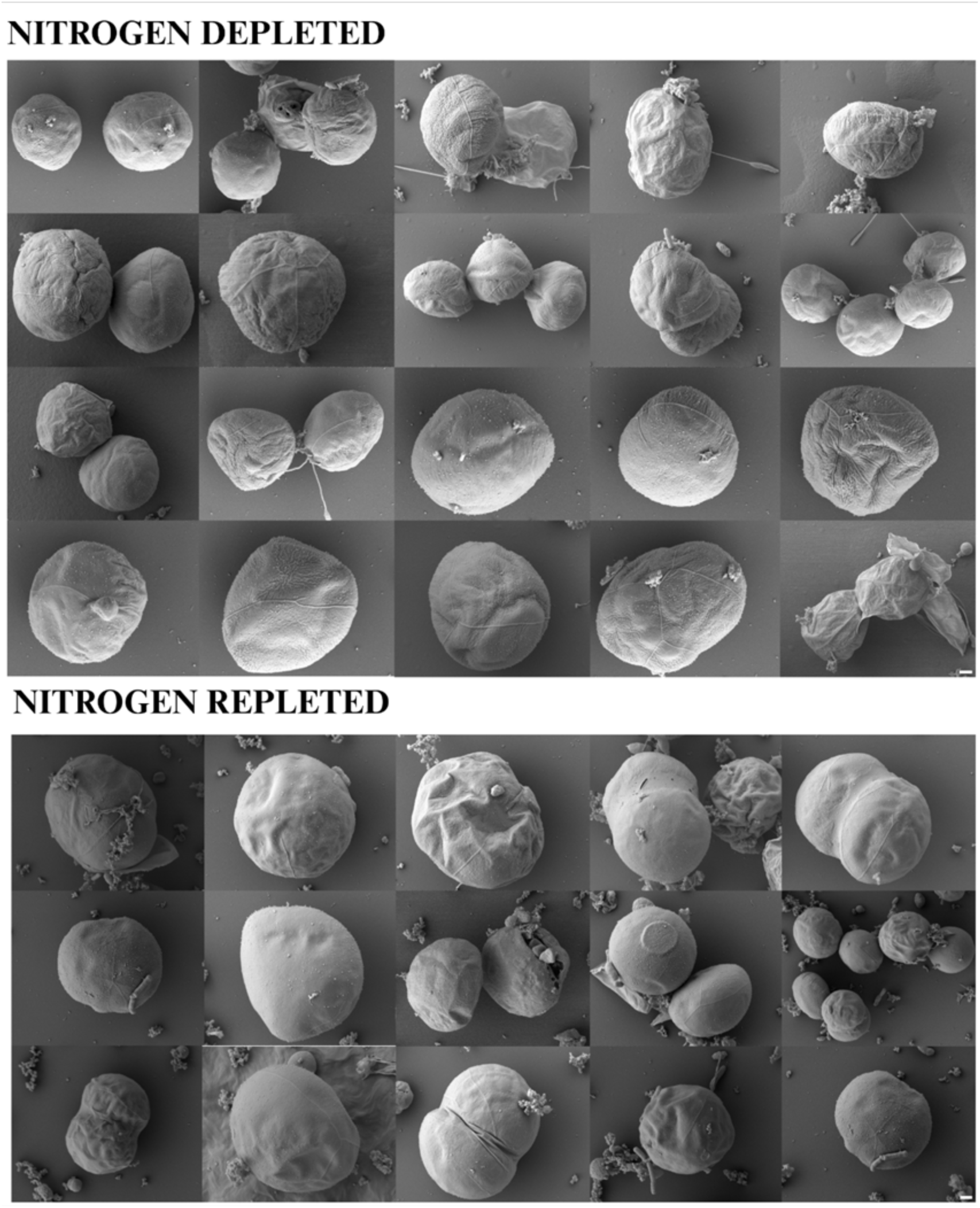
Scanning Electron micrographs of *Chromera velia* cells cultivated in nitrogen depleted and enriched conditions for 7 days. Scale bar = 1 µm.

**Figure S5.**
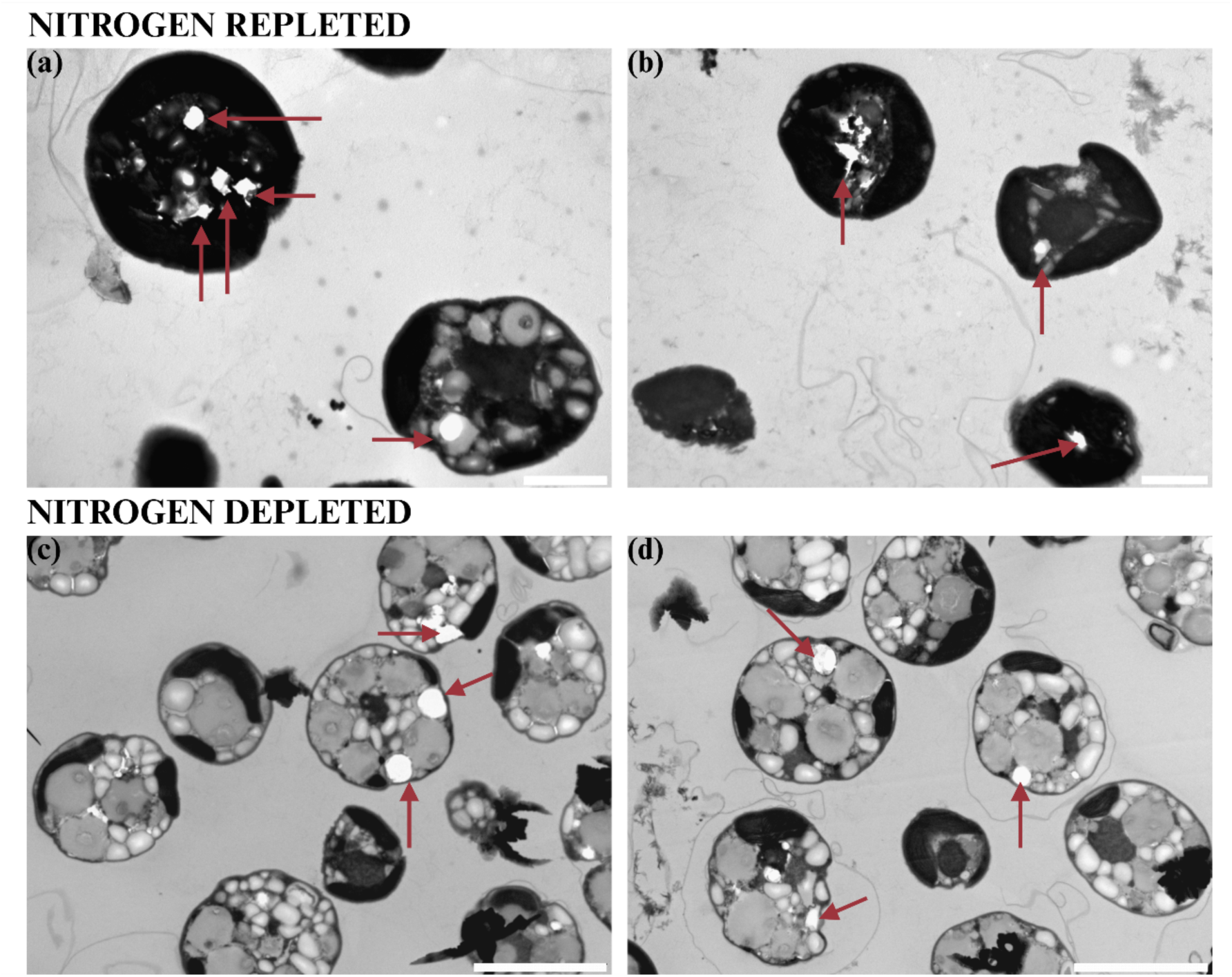
Transmission electron micrographs of *C. velia* cells. In standard ultra-thin sections, crystal containing vacuoles as well as starch grains often drop out of the sections (holes are marked with red arrows) and thus escape the attention of the viewer. Scale bars (a, b) = 2 µm, (c, d) = 5 µm.

